# IFN-γ-independent control of *M. tuberculosis* requires CD4 T cell-derived GM-CSF and activation of HIF-1α

**DOI:** 10.1101/2021.12.16.473015

**Authors:** Erik Van Dis, Huntly M Morrison, Daniel M Fines, Janet Peace Babirye, Lily H McCann, Sagar Rawal, Jeffery S Cox, Sarah A Stanley

## Abstract

The prevailing model of protective immunity to tuberculosis is that CD4 T cells produce the cytokine IFN-γ to activate bactericidal mechanisms in infected macrophages. Recent evidence has expanded this model, and it is now clear that CD4 T cells can control *M. tuberculosis* infection in the absence of IFN-γ production. To identify factors and pathways involved in IFN-γ-independent control, we developed a co-culture model using CD4 T cells isolated from the lungs of infected mice and *M. tuberculosis-infected* murine bone marrow-derived macrophages (BMDMs). We show that IFN-γ-independent control is primarily mediated by CD4 T cell production of the cytokine GM-CSF and requires activation of the macrophage transcription factor HIF-1α. HIF-1α activation drives a metabolic shift toward aerobic glycolysis and leads to the production of lipid droplets, both of which support host defense against infection. Surprisingly, recombinant GM-CSF is insufficient to rescue the absence of control by GM-CSF-deficient CD4 T cells during co-culture with BMDMs. In peritoneal macrophages, GM-CSF is sufficient to control growth, induces lipid droplet biogenesis, and requires HIF-1α expression for control. While HIF-1α-mediated control following IFN-γ stimulation requires nitric oxide, we find that HIF-1α activation by CD4 T cells and recombinant GM-CSF is nitric oxide-independent, implying a distinct pathway of activation. In addition to GM-CSF, CD4 T cells produce a factor that helps maintain phagosome membrane integrity during infection and blocks bacterial access to host lipids, a primary nutrient source. These results advance our understanding of CD4 T cell-mediated immunity to *M. tuberculosis*, clarify the role of nitric oxide as primarily immunomodulatory during *M. tuberculosis* infection, and reveal a novel mechanism for the activation of HIF-1α. Furthermore, we establish a previously unknown functional link between GM-CSF and HIF-1α and provide evidence that CD4 T cell-derived GM-CSF is a potent bactericidal effector.

## Introduction

Research into host immunity to *Mycobacterium tuberculosis* infection has the potential to improve the lives of billions of people around the world, yet major features of the immune response to tuberculosis (TB) remain poorly understood. Broadly, immunity to TB begins with an innate response that induces inflammation and the recruitment of phagocytes, followed by an adaptive response necessary to control infection. A critical aspect of this adaptive immune response is the activation and proliferation of *M. tuberculosis* specific CD4 T cells. Mice deficient in CD4 T cells are highly susceptible to TB, and the loss of CD4 T cells in patients suffering from AIDS is strongly correlated with re-activation of dormant *M. tuberculosis* infection (1, 2). The cytokine interferon (IFN)-γ is also required for control of TB. Mice deficient for IFN-γ signaling are among the most susceptible strains to *M. tuberculosis* infection, and recombinant IFN-γ activates the bactericidal capacity of macrophages (3–6). The known importance of CD4 T cells and IFN-γ has led to an enduring tenet of TB immunity: that CD4 T cells secrete IFN-γ to control *M. tuberculosis* growth in infected macrophages (7).

This basic understanding of protective immunity to TB has come under increasing scrutiny. Comparing different routes of inoculation with the vaccine strain *M. bovis* bacille Calmette–Guérin in mice shows that the frequency of IFN-γ-secreting CD4 T cells correlates more closely with disease severity than with protection from TB disease after *M. tuberculosis* challenge (8), and human trials with the vaccine candidate MVA85A showed that although this vaccine elicits significant numbers of IFN-γ-secreting CD4 T cells it does not lead to enhanced protection against infection (9, 10). Furthermore, while CD4 T cells are clearly important for TB control in humans, inherited mutations in components of the IFN-γ signaling pathway are not generally associated with susceptibility to *M. tuberculosis* but rather with enhanced susceptibility to non-tuberculosis mycobacteria such as *M. chelonae, M. smegmatis*, and *M. scrofulaceum* (11).

Basic research into immunity to TB has corroborated and expanded upon these findings. Adoptive transfer experiments in mice show that IFN-γ-deficient CD4 T cells can control *M. tuberculosis in vivo* (1, 12, 13) with IFN-γ production accounting for only 30% of CD4 T cell-mediated control in the lungs (13). Collectively, these findings point to an important unknown mechanism of control mediated by CD4 T cells that is independent of IFN-γ.

Immune resistance to TB requires a fine balance between pro-inflammatory effectors that control bacterial replication and anti-inflammatory immune regulation that prevents immunopathology. CD4 T cells themselves contribute to both of these arms, secreting pro-inflammatory molecules such as IFN-γ and TNFα and anti-inflammatory cytokines including IL-10 and TGF-β, complicating the interpretation of adoptive transfer experiments. Whether control by IFN-γ-deficient CD4 T cells following adoptive transfer is due to immunoregulatory effects or to the antibacterial effects of a specific IFN-γ-independent effector remains an open question. Thus far, adoptive transfer experiments have precluded a role for CD4 T cell expression of TNFα, Fas and perforin in IFN-γ-independent control (12), and have not demonstrated that a CD4 T cell-derived IFN-γ-independent effector can stimulate cell-intrinsic control of bacterial replication.

Recently, the cytokine granulocyte-macrophage colony-stimulating factor (GM-CSF) was shown to have a potential role in CD4 T cell-mediated control of TB. Recombinant GM-CSF (rGM-CSF) controls *M. tuberculosis* growth in mouse peritoneal macrophages and human monocytes, and GM-CSF-deficient (*Csf2^-/-^*) mice have significantly higher bacterial burden in the lungs compared to wild-type and succumb more rapidly to infection (14–16). However, *Csf2^-/-^* mice and humans with inherited mutations in the GM-CSF receptor have a defect in alveolar macrophage development which complicates the interpretation of whole-animal knockout experiments (17, 18). Finally, adoptive transfer experiments show that *Csf2^-/-^* CD4 T cells only exhibit less control than wild-type when transferred into mice deficient for GM-CSF (15), so it remains unclear whether CD4 T cell production of GM-CSF induces macrophage-intrinsic control of *M. tuberculosis*.

In this study, we use an *in vitro* co-culture system to determine whether an IFN-γ-independent CD4 T cell effector can induce cell intrinsic control of *M. tuberculosis* replication. We show that lung-derived CD4 T cells and multiple *in vitro*-differentiated T cell subsets exhibit IFN-γ-independent control in infected macrophages via a secreted and proteinaceous effector, and we use RNA sequencing and cytokine profiling to investigate possible mechanisms. Like IFN-γ-mediated control, IFN-γ-independent control by CD4 T cells requires macrophage expression of the transcription factor hypoxia inducible factor-1α (HIF-1α), activation of which leads to a metabolic switch to aerobic glycolysis and the formation of macrophage lipid droplets to control bacterial growth. These changes occur independent of the IFN-γ-induced second messenger nitric oxide (NO)—normally required for HIF-1α activation during *M. tuberculosis* infection— indicating a novel mechanism of HIF-1α activation. Furthermore, we show that while there is no role for CD4 T cell production of TNFα, Type I IFN, CD153 (*Tnfsf8*) or CD40, CD4 T cell secretion of GM-CSF is required for IFN-γ-independent control. Recombinant GM-CSF is sufficient to control *M. tuberculosis* in mouse peritoneal macrophages, requires HIF-1α expression for control and correlates with the production of macrophage lipid droplets. However, GM-CSF is insufficient for control in bone marrow-derived macrophages (BMDMs) and, surprisingly, does not rescue the loss of IFN-γ-independent control by GM-CSF-deficient CD4 T cells, indicating either differences in recombinant and CD4 T cell-derived GM-CSF function or an unknown second CD4 T cell effector necessary for GM-CSF-mediated control in BMDMs. These results advance our understanding of CD4 T cell-mediated control of *M. tuberculosis*, establish CD4 T cell-derived GM-CSF as a potent bactericidal effector and, by uncovering a novel mechanism for HIF-1α activation, emphasize the central importance of HIF-1α for intracellular immunity to TB.

## Results

### Lung-derived and *in vitro*-differentiated CD4 T cells exhibit IFN-γ-independent control of *M. tuberculosis*

To confirm that CD4 T cells can control *M. tuberculosis* independent of IFN-γ, we cultured CD4 T cells derived from the lungs of infected mice with infected *Ifngr^-/-^* BMDMs and observed dose-dependent IFN-γ-independent control of bacterial growth (Fig 1A, S1A).

**Fig 1.**
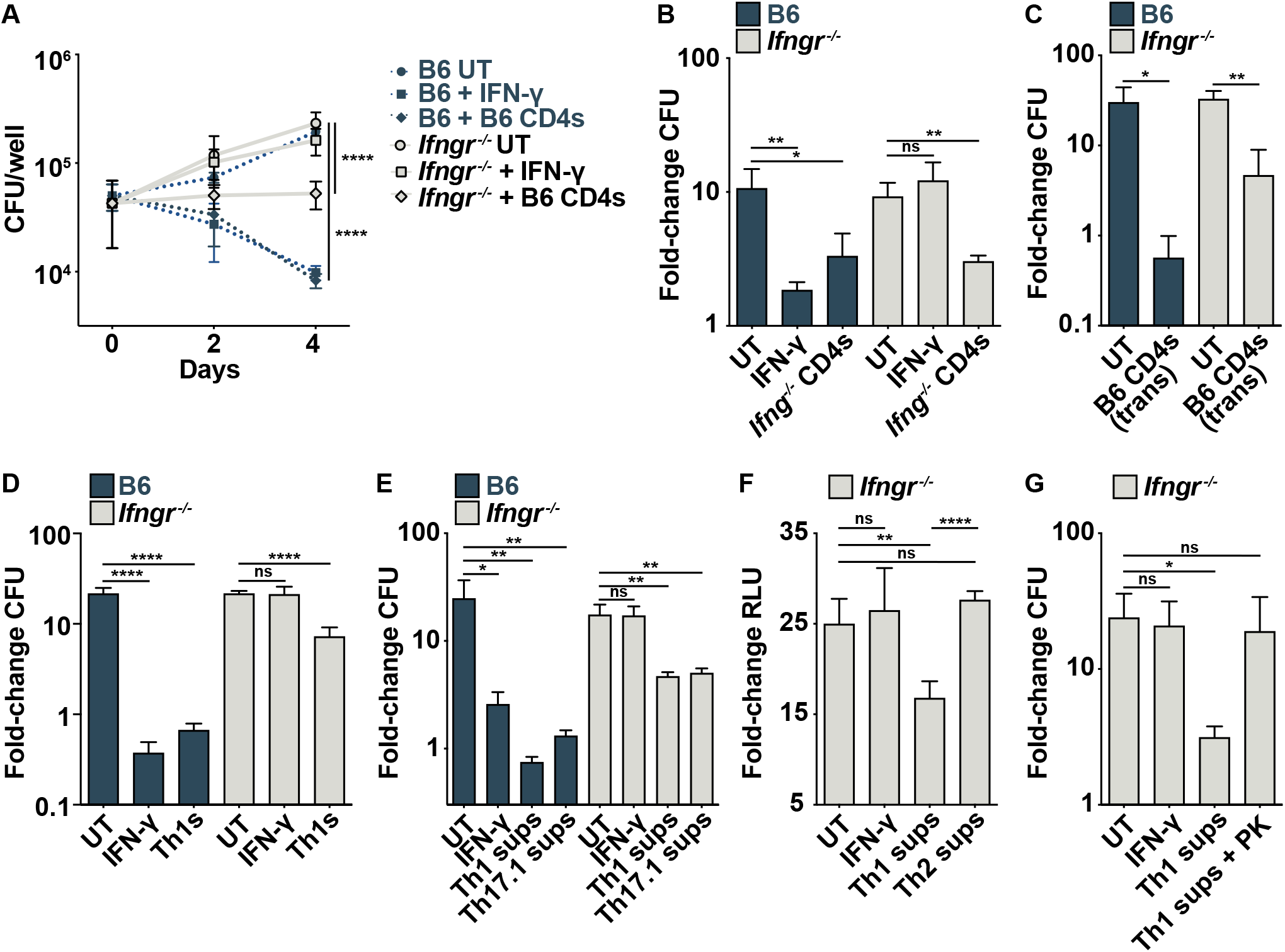
Lung-derived and in vitro-differentiated CD4 T cells control M. tuberculosis growth independent of IFN-γ. (**A**) CFU/well at d 0, 2 and 4 postinfection for wild-type and *Ifngr^-/-^* BMDMs co-cultured with lung-derived wild-type CD4 T cells. (**B**) CFU fold-change at d 5 postinfection for wild-type and *Ifngr^-/-^* BMDMs co-cultured with lung-derived *Ifng^-/-^* CD4 T cells. (**C**) CFU fold-change at d 4 postinfection for wild-type and *Ifngr^-/-^* BMDMs cultured with wild-type lung-derived CD4 T cells in transwells. (**D**) CFU fold-change at d 5 postinfection for wild-type and *Ifngr^-/-^* BMDMs co-cultured with *in vitro* differentiated Th1 cells. (**E**) CFU fold-change at d 4 postinfection for wild-type and *Ifngr^-/-^* BMDMs treated with Th1 or Th17.1 supernatants (sups). (**F**) RLU fold-change at d 5 postinfection for *Ifngr^-/-^* BMDMs treated with Th1 or Th2 supernatants. (**G**) CFU fold-change at d 5 postinfection for *Ifngr^-/-^* BMDMs treated with Th1 supernatants +/- Proteinase K (PK). Figures are representative of two (G) or three or more (A)-(F) experiments. Error bars are SD from four replicate wells (A)-(B), (D)-(G) or three replicate wells (C),*p<0.05, **p<0.01, ****p<0.0001 by unpaired t-test.

Similarly, CD4 T cells from the lungs of infected *Ifng^-/-^* mice controlled bacterial growth during co-culture with wild-type BMDMs (Fig 1B). Conditioned media from CD4 T cells and CD4 T cells in transwells both controlled growth in *Ifngr^-/-^* BMDMs showing that IFN-γ-independent control is mediated by a secreted effector (Fig 1C, S1B). To ask whether a specific T cell subset is sufficient for IFN-γ-independent control, we *in vitro* differentiated Th1, Th2 and Th17.1 T cells using *M. tuberculosis* specific TCR transgenic mice. Th1s restricted bacterial growth during co-culture with *Ifngr^-/-^* BMDMs (Fig 1D), and supernatants from Th1s exhibited dose-dependent IFN-γ-independent control (Fig 1E, S1C, S1D). Supernatants from the closely related Th17.1 T cell subset also demonstrated IFN-γ-independent control (Fig 1E), while supernatants from Th2s did not (Fig 1F). Taken together, these results show that lung-derived CD4 T cells and *in vitro* differentiated Th1s and Th17.1s secrete an IFN-γ-independent effector that induces cell-intrinsic control of *M. tuberculosis*. To test whether this effector is amino acid-based, we treated Th1 supernatants with proteinase K and observed a loss of IFN-γ-independent control (Fig 1G). Thus, a proteinaceous effector secreted by CD4 T cells activates BMDMs to control *M. tuberculosis* independent of IFN-γ.

### IFN-γ-independent control requires an NO-independent HIF-1α-mediated shift to aerobic glycolysis

To identify macrophage signaling pathways induced by CD4 T cells independent of IFN-γ, we performed RNA sequencing on *M. tuberculosis-infected* wild-type and *Ifngr^-/-^* BMDMs after 24 hours of co-culture with lung CD4 T cells. CD4 T cells induced differential regulation of >1800 genes in wild-type BMDMs and >800 genes in *Ifngr^-/-^* BMDMs compared to untreated (Fig 2A, 2B). Of these, 192 genes were altered in an IFN-γ-independent manner, with differential expression in both wild-type and *Ifngr^-/-^* BMDMs during CD4 co-culture (Fig 2A). Using transcripts with >2-fold upregulation in both wild-type and *Ifngr^-/-^* BMDMs, gene ontology (GO) analysis revealed cytokine-mediated signaling, positive regulation of cytokine production, and inflammatory responses as the top three enriched GO terms following co-culture with CD4 T cells (19–21). During *M. tuberculosis* infection of macrophages, IFN-γ signaling causes a metabolic switch to aerobic glycolysis that is required for IFN-γ-mediated control (22).

**Fig 2.**
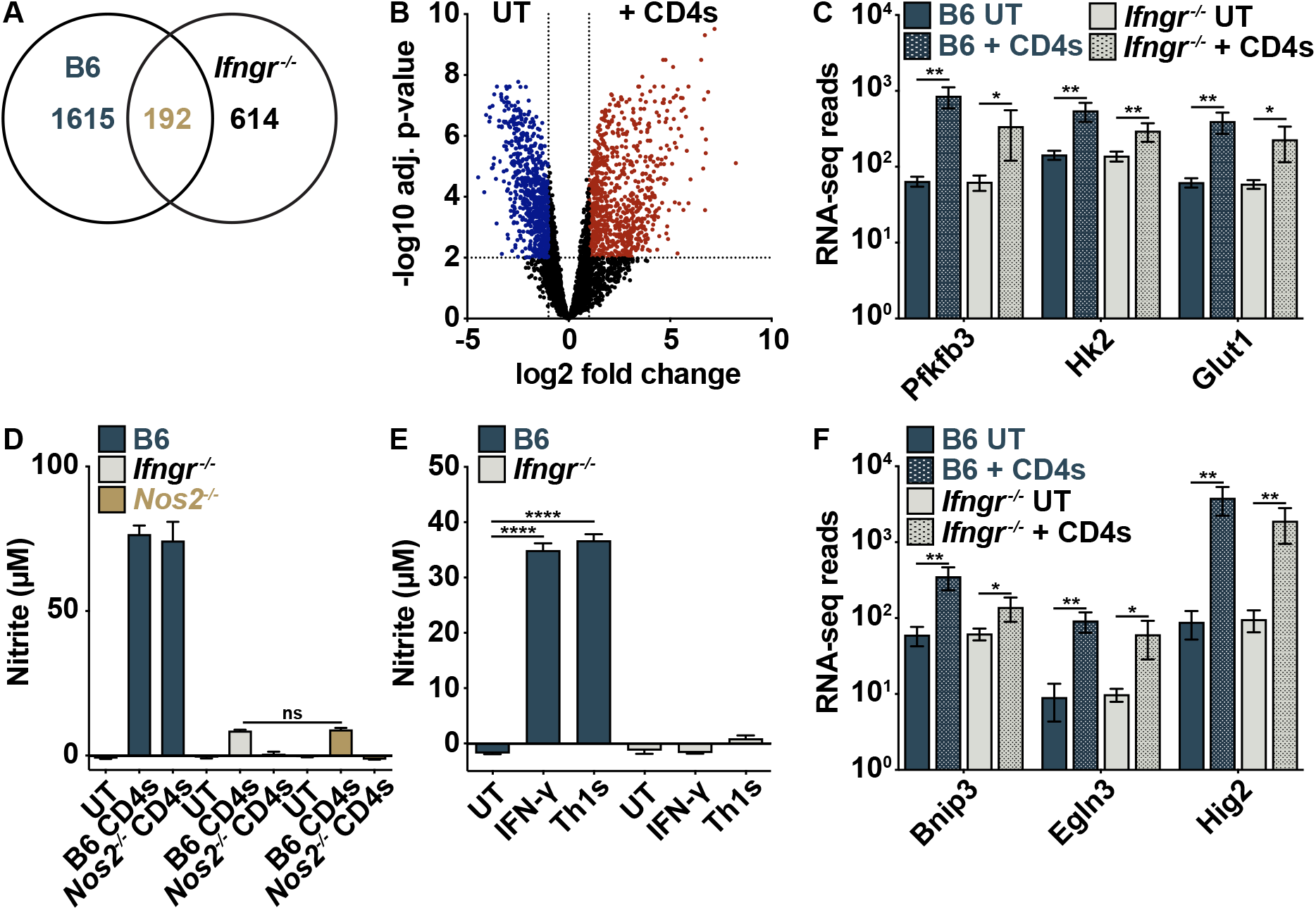
CD4 T cells induce an IFN-γ- and NO-independent increase in glycolytic gene expression. (**A**)-(**B**) RNA-seq at 24 h postinfection for wild-type and *Ifngr^-/-^* BMDMs co-cultured with wild-type lung CD4 T cells. (A) Venn diagram showing overlap of CD4 T cell-regulated genes in wild-type and *Ifngr^-/-^* BMDMs with at least 2-fold change in gene expression and adj. p-value >0.05 relative to UT. (B) Volcano plot of genes in *Ifngr^-/-^* BMDMs showing >2-fold change in gene expression and adj. p-value >0.05 between UT and CD4 T cell co-culture. (**C**) RNA-seq reads of glycolytic genes. (**D**) Griess assay at 48 h postinfection for wild-type, *Ifngr^-/-^*, and *Nos2^-/-^* BMDMs co-cultured with a 10:1 ratio of wild-type or *Nos2^-/-^* lung-derived CD4 T cells. (**E**) Griess assay at 48 h postinfection for wild-type and *Ifngr^-/-^* BMDMs co-cultured with Th1 cells. (**F**) RNA-seq reads of HIF-1α target genes. Figures are representative of two (E) or three (D) independent experiments. Error bars are SD from three independent experiments (C), (F) or four replicate wells (D)-(E), *p<0.05, **p<0.01, ****p<0.0001 by unpaired t-test.

Interestingly, CD4 T cells also induced significant upregulation of genes associated with aerobic glycolysis and an increase in glucose uptake, both with and without IFN-γ signaling (Fig 2C, S2A). Nitric oxide (NO) production by inducible nitric oxide synthase (iNOS) is required for the IFN-γ-mediated shift to aerobic glycolysis, and iNOS-deficient (*Nos2^-/-^*) BMDMs have a defect in cell intrinsic control of *M. tuberculosis* following IFN-γ signaling (23). To explore a role for NO in IFN-γ-independent control, we first asked whether NO is produced during CD4 T cell co-culture. While NO is produced in abundance by IFN-γ-activated BMDMs during *M. tuberculosis* infection, there is little to no NO production induced by CD4 T cells or Th1 supernatants in the absence of IFN-γ signaling (Fig 2D, 2E). A low level of NO was detected during CD4 T cell co-cultures only when CD4 T cells themselves expressed iNOS, and this amount of CD4 T cell-derived NO does not contribute to control of bacterial growth in macrophages (Fig 2D, S2B). Therefore, IFN-γ-independent control by lung CD4 T cells and *in vitro* differentiated Th1s is independent of NO.

IFN-γ signaling stabilizes and activates the transcription factor HIF-1α during *M. tuberculosis* infection (22). HIF-1α is required for the IFN-γ-mediated metabolic switch to aerobic glycolysis, and HIF-1α-deficient (*Hif1a^-/-^*) BMDMs have a defect in IFN-γ-mediated control (22). NO helps stabilize HIF-1α and the two form a positive feedback loop that is necessary for the antibacterial effect of IFN-γ (23). Surprisingly, despite an absence of NO and IFN-γ signaling, we observed significant upregulation of multiple HIF-1α target genes in *Ifngr^-/-^* BMDMs during CD4 T cell co-culture (Fig 2F), and bioinformatic analysis revealed an overrepresentation of genes with predicted HIF-1α binding sites upregulated in *Ifngr^-/-^* BMDMs (Fig S2C).

To explore a role for HIF-1α in IFN-γ-independent control, we treated *Ifngr^-/-^* BMDMs with Th1 supernatants and observed stabilization of HIF-1α by western blot (Fig 3A). As has been demonstrated for IFN-γ-mediated HIF-1α stabilization (22), this was dependent on a metabolic switch to aerobic glycolysis since concurrent treatment of Th1 supernatants with the glycolysis inhibitor 2-deoxy-D-glucose (2-DG) abolished HIF-1α stabilization (Fig 3A). Importantly, *Ifng^-/-^* CD4 T cells did not control bacterial growth in *Hif1a^-^* BMDMs (Fig 3B) and 2-DG treatment partially reduced IFN-γ-independent control by Th1 supernatants (Fig S3A). These results demonstrate that HIF-1α stabilization and aerobic glycolysis are necessary for IFN-γ-independent control of *M. tuberculosis* by CD4 T cells. Activation of HIF-1α mediates lipid droplet (LD) biogenesis in *M. tuberculosis* infected BMDMs, which supports host immunity by serving as a platform for eicosanoid production (24). We observed a similar increase in the number and size of LDs between infected wild-type and *Ifngr^-/-^* BMDMs after treatment with Th1 supernatants (Fig 3C-G, S3B). The observation that HIF-1α activation and LD accumulation occur independent of IFN-γ and NO during *M. tuberculosis* infection, and that HIF-1α is required for IFN-γ-independent control, solidifies HIF-1α as a critical signaling node after CD4 T cell activation of infected macrophages and suggests that pathways of IFN-γ-dependent and IFN-γ-independent control by CD4 T cells converge on HIF-1α activation.

**Fig 3.**
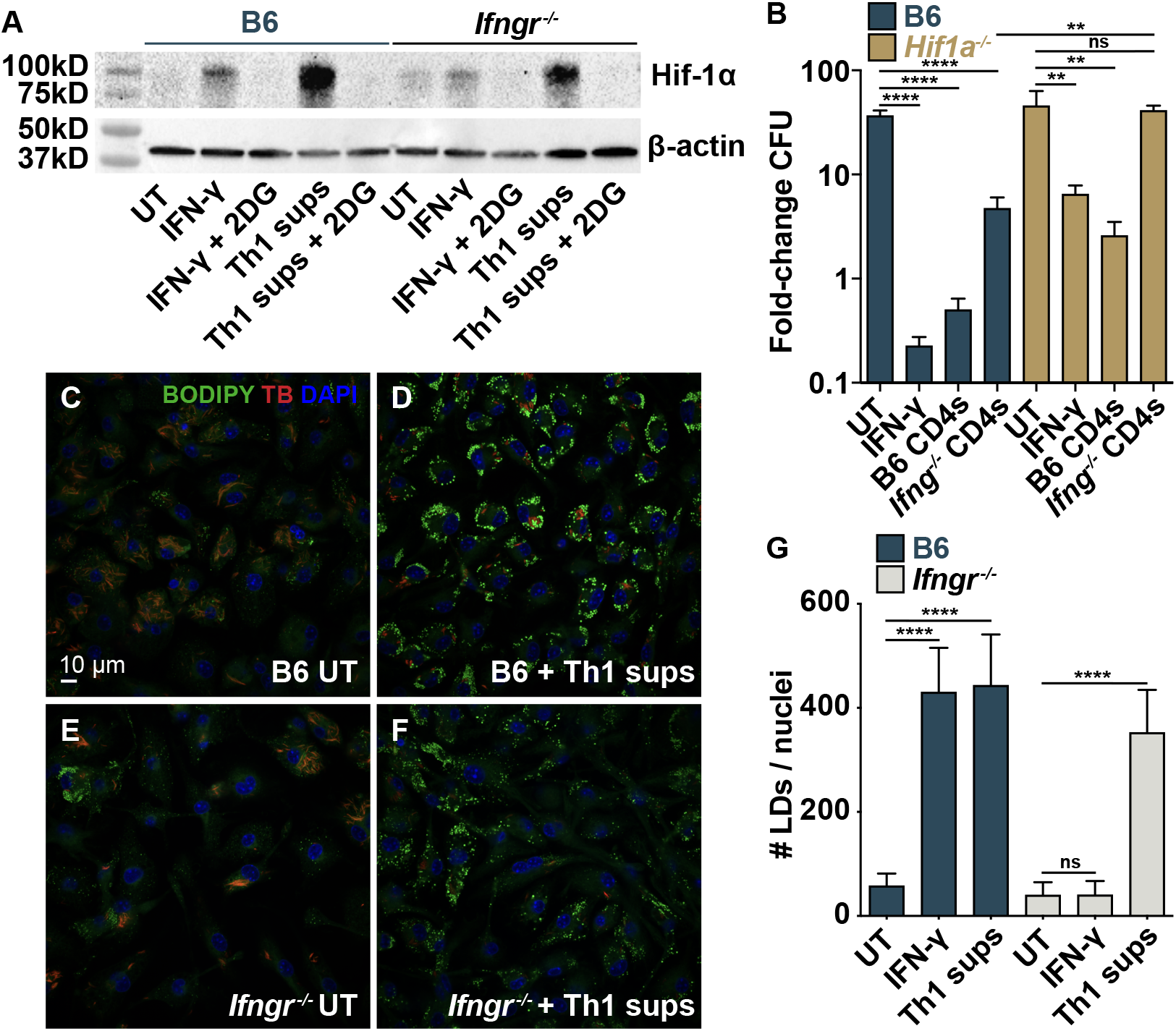
IFN-γ-independent control requires the transcription factor HIF-1α. (**A**) Western blot for HIF-1α on cell lysates 12 h postinfection for wild-type and *Ifngr^-/-^* BMDMs treated with Th1 supernatants and 2-DG. (**B**) CFU fold-change at d 5 postinfection for wild-type and *Hif1a^-/-^* BMDMs co-cultured with wild-type or *Ifng^-/-^* lung-derived CD4 T cells. (**C**)-(**F**) Microscopy for host lipid droplets (LDs) at d 3 postinfection for wild-type and *Ifngr^-/-^* BMDMs treated with Th1 supernatants. (**G**) Quantification of (C)-(F) for average number of LDs per BMDM. Figures are representative of two (A) or three (B) independent experiments or 48 images from four replicate wells (G), **p<0.01, ****p<0.0001 by unpaired t-test.

### CD4 T cell-derived GM-CSF is necessary for IFN-γ-independent control of *M. tuberculosis*

To determine which secreted effector is necessary for IFN-γ-independent control, we performed cytokine profiling of Th1 and Th17.1 supernatants (Fig 4A). Multiple inflammatory cytokines previously implicated in immune control of *M. tuberculosis* were identified, including TNFα, IFN-α, IL-1β, IFN-γ, and GM-CSF. *Ifng^-/-^* CD4 T cells had no loss of control in co-culture with *Tnfr1^-/-^/Tnfr2^-/-^* BMDMs compared to wild-type (Fig 4B), and Th1 supernatants had no loss of control in *Ifnar^-/-^/Ifngr^-/-^* BMDMs compared to *Ifngr^-/-^* BMDMs (Fig 4C). Additionally, blocking antibodies for CD40 had no effect on IFN-γ-independent control by lung CD4 T cells (Fig S4A). Thus, IFN-γ-independent CD4 T cell control of *M. tuberculosis* in BMDMs does not require TNFα, Type I IFN, or CD40. The CD4 T cell surface molecule CD30 ligand (*Tnfsf8*) has been implicated in IFN-γ-independent control of *M. tuberculosis in vivo* (25). However, we observed little to no expression of the gene for CD30 (*Tnfrsf8*) in BMDMs in the presence or absence of CD4 T cells (Fig S4B) and no loss of IFN-γ-independent control in BMDMs doubly deficient for CD30 and IFNγR (*Tnfrsf8^-/-^/Ifngr^-/-^*) compared to *Ifngr^-/-^* BMDMs alone (Fig 4D). CD30 and CD30 ligand are cell-surface molecules so, together with the observation that IFN-γ-independent control does not require cell-to-cell contact (Fig 1C, S1B), these results argue that the role of CD30 ligand on CD4 T cells during TB infection is likely immunoregulatory rather than directly bactericidal.

**Fig 4.**
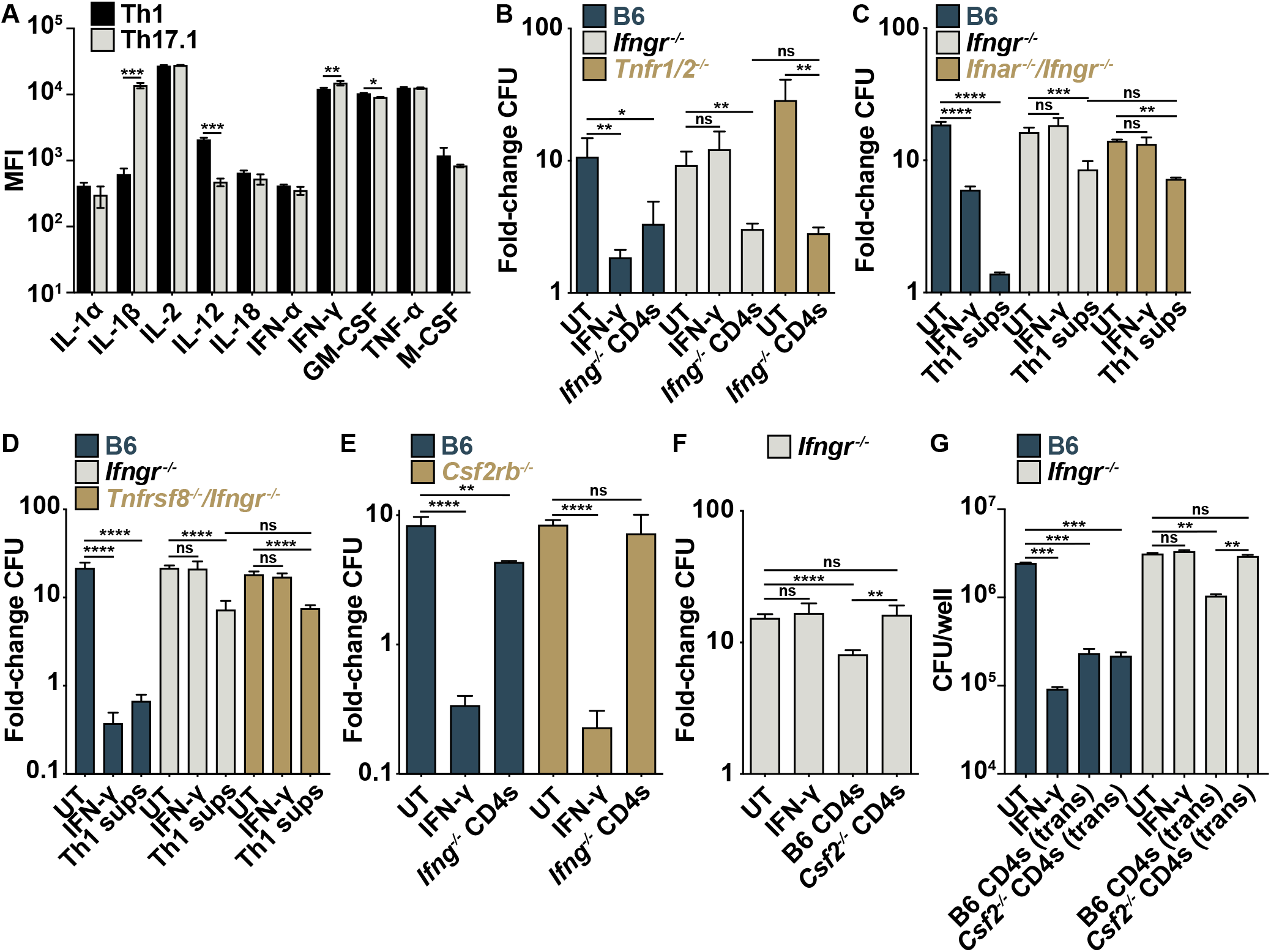
IFN-γ-independent control of M. tuberculosis requires CD4 T cell-derived GM-CSF. (**A**) Mean fluorescence intensity (MFI) of select cytokines in *in vitro* differentiated Th1 and Th17.1 supernatants. (**B**) CFU fold-change at d 5 postinfection for wild-type, *Ifngr^-/-^*, and *Tnfr1/2^-/-^* BMDMs co-cultured with *Ifng^-/-^* lung-derived CD4 T cells. (**C**) CFU fold-change at d 5 postinfection for wild-type, *Ifngr^-/-^*, and *Ifnar^-/-^/Ifngr^-/-^* BMDMs treated with Th1 supernatants. (**D**) CFU fold-change at d 5 postinfection for wild-type, *Ifngr^-/-^* and *Tnfrsf8^-/-^/Ifngr^-/-^* double-knockout BMDMs treated with Th1 supernatants. (**E**) CFU fold-change at d 5 postinfection for wild-type and *Csf2rb^-/-^* BMDMs co-cultured with *Ifng^-/-^* lung-derived CD4 T cells. (**F**) CFU fold-change at d 5 postinfection for *Ifngr^-/-^* BMDMs co-cultured with wild-type or *Csf2^-/-^* lung-derived CD4 T cells. (**G**) CFU-fold change at d 4 postinfection for wild-type and *Ifngr^-/-^* BMDMs cultured with wild-type or *Csf2^-/-^* lung-derived CD4 T cells in transwells. Figures represent seven (Th1) or three (Th17.1) biological replicates (A) or are representative of two (B), (D), (G) or at least three (C), (F) independent experiments. Error bars are SD from biological replicates (A), four replicate wells (B)-(F) or three replicate wells (G), *p<0.05, **p<0.01, ***p<0.001, ****p<0.0001 by unpaired t-test.

We next tested a role for the cytokine GM-CSF, which is secreted by Th1 and Th17.1 T cells (Fig 4A) and can be found in the supernatant of BMDMs co-cultured with CD4 T cells from the lungs of wild-type and *Ifng^-/-^* mice (Fig S4C). *Csf2ra* and *Csf2rb*, genes for the two GM-CSF receptor subunits, are expressed in BMDMs and transcription increases more than 3-fold following CD4 co-culture (Fig S4D-E). Importantly, while *Ifng^-/-^* CD4s control *M. tuberculosis* in wild-type BMDMs, this control is lost in BMDMs lacking the GM-CSF receptor (*Csf2rb^-/-^*) (Fig 4E). Furthermore, wild-type but not GM-CSF deficient (*Csf2^-/-^*) CD4 T cells control bacterial growth during co-culture or when cultured in transwells with *Ifngr^-/-^* BMDMs (Fig 4F-G). Taken together, these results show that lung CD4 T cells secrete GM-CSF to control *M. tuberculosis* growth in macrophages independent of IFN-γ.

### GM-CSF activates HIF-1α to restrict *M. tuberculosis* in peritoneal macrophages

As reported, recombinant GM-CSF (rGM-CSF) is sufficient to control *M. tuberculosis* growth in peritoneal macrophages (Fig 5A) (15, 26). However, despite a clear requirement for GM-CSF signaling in BMDMs during CD4 T cell co-culture, rGM-CSF is not sufficient to induce control of *M. tuberculosis* in BMDMs (Fig 5A), regardless of the biological source of recombinant protein (Fig S5A). Moreover, rGM-CSF from multiple sources did not rescue the loss of control by *Csf2^-/-^* CD4 T cells in co-culture with *Ifngr^-/-^* BMDMs (Fig 5B, S5B). Thus, while GM-CSF is sufficient to control *M. tuberculosis* in peritoneal macrophages it is necessary but not sufficient for CD4 T cell-mediated control in BMDMs, reflecting either differences in the ability of CD4 T cell-derived and recombinant GM-CSF to effectively signal to BMDMs or secondary defects in *Csf2^-/-^* CD4 T cells required for IFN-γ-independent control.

**Fig 5.**
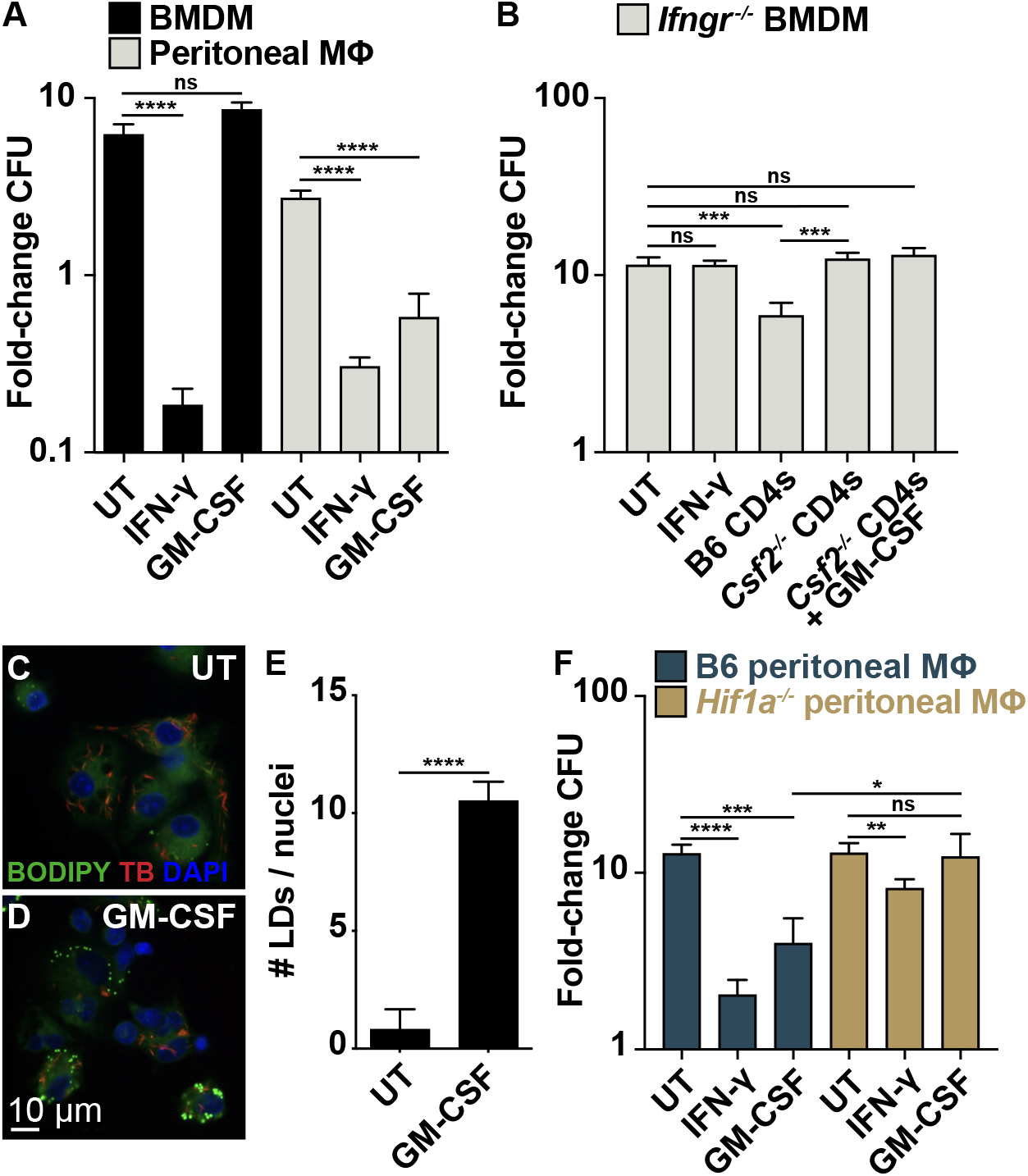
GM-CSF activates HIF-1α to restrict M. tuberculosis in peritoneal macrophages. (**A**) CFU fold-change at d 4 postinfection for BMDMs and peritoneal macrophages treated with GM-CSF. (**B**) CFU fold-change at d 5 postinfection for *Ifngr^-/-^* BMDMs co-cultured with wild-type or *Csf2^-/-^* lung CD4 T cells and GM-CSF. (**C**)-(**D**) Microscopy for host lipid droplets (LDs) at d 3 postinfection for wild-type peritoneal macrophages treated with GM-CSF. (**E**) Quantification of (C)-(D) for average number of LDs per peritoneal macrophage treated with GM-CSF. (**F**) CFU fold-change at d 4 postinfection for wild-type and *Hif1a^-/-^* peritoneal macrophages treated with GM-CSF. All figures are representative of three or more experiments. Error bars are SD from four replicate wells (A)-(B), (F) or 48 images from four replicate wells (E), *p<0.05, **p<0.01, ***p<0.001, ****p<0.0001 by unpaired t-test.

The antibacterial mechanisms downstream of GM-CSF signaling are still unclear. As expected, treatment of *M. tuberculosis-infected* peritoneal macrophages with rGM-CSF did not induce production of NO (Fig S5C). Since IFN-γ-independent control of *M. tuberculosis* by lung CD4 T cells is NO-independent and requires HIF-1α (Fig 3C), we tested a role for HIF-1α in GM-CSF-mediated control in peritoneal macrophages. rGM-CSF treatment leads to an increase in the number and size of LDs which is a prominent effect of HIF-1α activation (Fig 5C-E, S5D), and peritoneal macrophages lacking HIF-1α have a significant defect in GM-CSF-mediated control compared to wild-type (Fig 5F). Activation of HIF-1α in the absence of NO indicates that the mechanism of HIF-1α activation following GM-CSF signaling is distinct from the NO-mediated HIF-1α stabilization initiated by IFN-γ.

### CD4 T cells reduces cytosolic access of *M. tuberculosis* independent of IFN-γ and GM-CSF

Autophagy is a major contributor to immune defense against *M. tuberculosis* in macrophages (27, 28). *M. tuberculosis* is targeted by autophagy following permeabilization of the phagosome by the ESX-1 secretion system, which the bacteria use to gain access to the cytosol (27–29). Early autophagic targeting of *M. tuberculosis* involves deposition of ubiquitin chains around bacteria via Parkin and the cGAS-STING pathway (30, 31). We hypothesized that CD4 T cells may control bacterial growth in an IFN-γ-independent manner by inducing autophagy, leading to an increase in ubiquitin colocalization with intracellular bacteria. We observed, however, that treatment of infected macrophages with Th1 supernatants leads to a decrease in ubiquitin recruitment to *M. tuberculosis* in an IFN-γ-independent manner (Fig 6A), which may be caused by reduced phagosome perforation and bacterial access to the cytosol. In support of this, treatment with Th1 supernatants also results in a significant reduction in the ability of *M. tuberculosis* to accumulate intracellular lipid inclusions, which serve as an important host-derived carbon source for the bacteria (Fig 6B) (32, 33).

**Fig 6.**
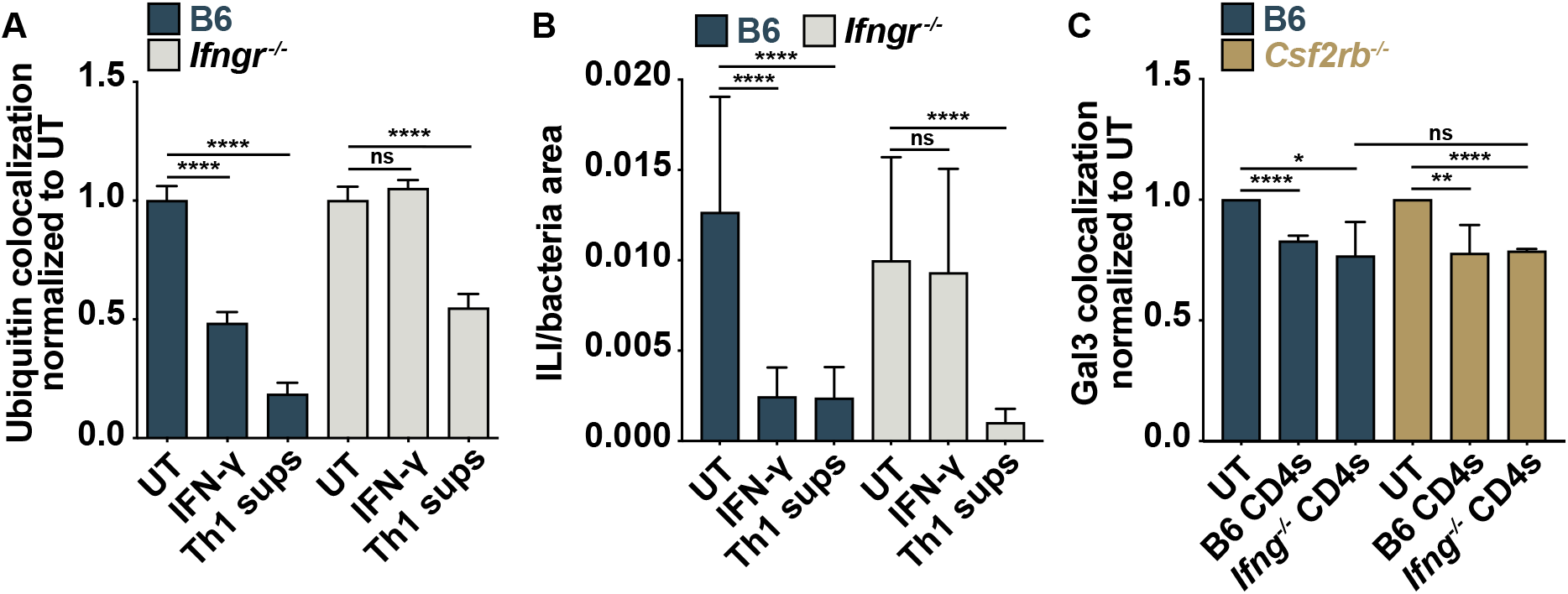
CD4 T cells reduces cytosolic access of M. tuberculosis independent of IFN-γ and GM-CSF. (**A**) Percent colocalization between bacteria and ubiquitinylated proteins at 8 h postinfection for wild-type and *Ifngr^-/-^* BMDMs treated with Th1 supernatants, normalized to untreated for each BMDM genotype. (**B**) Microscopy for ILI intensity per bacteria area at d 3 postinfection for wild-type and *Ifngr^-/-^* BMDMs treated with Th1 supernatants. (**C**) Percent colocalization between bacteria and Galectin 3 (Gal3) at 8 h postinfection for wild-type and *Csf2rb^-/-^* BMDMs cultured with lung-derived wild-type or *Ifng^-/-^* CD4 T cells in transwells, normalized to untreated for each BMDM genotype. Error bars are SD from 48 images from four replicate wells (A)-(B) or from four replicate wells with 9 images per well (C), *p<0.05, **p<0.01, ****p<0.0001 by unpaired t-test.

To test a role for GM-CSF in this phenotype, we cultured lung-derived wild-type or *Csf2^-/-^* CD4 T cells in transwells over wild-type or *Ifngr^-/-^* BMDMs and performed microscopy for bacterial colocalization with Galectin 3 (Gal3), which forms puncta on damaged phagosomes and is a more direct measure of membrane integrity. Consistent with our findings using Th1 supernatants, both wild-type and *Ifng^-/-^* CD4 T cells lead to a significant reduction in Gal3-bacteria colocalization in wild-type BMDMs, demonstrating IFN-γ-independent CD4 T cell-induced renitence of *M. tuberculosis* containing phagosomes (Fig 6C). However, this reduction did not require GM-CSF, as there was no difference between the wild-type and *Csf2rb^-/-^* BMDMs. The connection between phagosome integrity, nutrient restriction, and IFN-γ-independent CD4 T cell control is an ongoing area of investigation.

## Discussion

Immunity to *M. tuberculosis* requires CD4 T cells and IFN-γ to activate effector functions within infected macrophages. This is supported by the extreme susceptibility of *Ifng^-/-^* and *Rag1^-/-^* mice, the high co-morbidity between TB and low CD4 T cell counts caused by AIDS, and the ability of recombinant IFN-γ to control bacterial growth in mouse and human macrophages *in vitro* (1–6, 34). However, while IFN-γ production by CD4 T cells is likely necessary for full control of infection, it has become clear that IFN-γ is a poor biomarker for effective CD4 T cell immunity to TB and that CD4 T cells are capable of controlling infection even in the absence of IFN-γ (1, 7, 35). Here, we use an *in vitro* CD4 T cell-macrophage co-culture model to confirm the finding that CD4 T cells can control *M. tuberculosis* in the absence of IFN-γ, and we demonstrate that this control is mediated by a secreted, antibacterial effector produced by lung-derived CD4 T cells and by *in vitro* differentiated Th1 and Th17.1 T cells.

RNA sequencing of *M. tuberculosis-infected* BMDMs after CD4 T cell co-culture revealed an IFN-γ-independent program of HIF-1α-driven transcription and significant upregulation of genes involved in glycolysis, reminiscent of the effects observed following IFN-γ activation. CD4 T cells also induced the development of macrophage LDs—a direct effect of HIF-1α stabilization—in the absence of IFN-γ signaling. LDs support macrophage defense against infection, and lung granulomas in TB patients are surrounded by “foamy macrophages” that contain large accumulations of LDs (36, 37). Importantly, we demonstrate that HIF-1α expression is required for IFN-γ-independent control and show that, unlike HIF-1α stabilization following IFN-γ signaling, CD4 T cells stabilize HIF-1α in *Ifngr^-/-^* macrophages in the absence of NO. These findings help clarify the role of NO during *M. tuberculosis* infection. While NO has long been thought to be directly bactericidal to *M. tuberculosis*, the regulatory role of NO as a second messenger has gained recent appreciation (38, 39), including work from our laboratory showing that HIF-1α and iNOS form a positive feedback loop required for IFN-γ-mediated control of *M. tuberculosis* in BMDMs (23). Furthermore, we have shown that HIF-1α and iNOS participate in the same pathway following IFN-γ activation of BMDMs (23). IFN-γ- and NO-independent HIF-1α stabilization during CD4 T cell co-culture positions NO upstream of HIF-1α following IFN-γ signaling and implicates NO as primarily a regulatory, rather than bactericidal, molecule during *M. tuberculosis* infection, at least in the context of CD4 T cell activation of macrophages. Furthermore, the need for HIF-1α expression to mediate both IFN-γ-mediated and IFN-γ-independent CD4 T cell control emphasizes the central role of HIF-1α in cell-intrinsic control of infection and indicates that these two pathways of macrophage activation converge on HIF-1α activation. Finally, we show that GM-CSF control of *M. tuberculosis* in peritoneal macrophages requires HIF-1α. This is the first demonstration of a functional link between GM-CSF and HIF-1α during *M. tuberculosis* infection, although this relationship has precedence in the literature in other contexts. In mouse neural progenitor cells GM-CSF treatment induces a PI3K–NF-κB signaling pathway that increases HIF-1α expression (40), and in a mouse melanoma model GM-CSF treatment induces macrophage secretion of vascular endothelial growth factor (VEGF) in a HIF-1α-dependent manner (41). Research into the signaling pathways that mediate NO-dependent and -independent HIF-1α stabilization will help elucidate mechanisms of cell-intrinsic control of *M. tuberculosis*.

Given the importance of autophagy for macrophage control of *M. tuberculosis*, we reasoned that increased autophagic targeting may contribute to IFN-γ-independent control by CD4 T cells. Instead, we observed a decrease in ubiquitin recruitment to *M. tuberculosis*-containing phagosomes, a decrease in lipid uptake by the bacteria, and a decrease in phagosome damage as measured by Gal3 puncta colocalization that was independent of both IFN-γ and GM-CSF. IFN-γ and LPS stimulation can induce resistance to phagosome membrane damage in macrophages that is important for host resistance to another intracellular pathogen, *Listeria monocytogenes* (42–44). Additionally, IFN-γ stimulation has been shown to help maintain phagosome integrity during *M. tuberculosis* infection in a Rab20-dependent manner (45). Our results indicate that CD4 T cells secrete an IFN-γ- and GM-CSF-independent effector that reinforces the integrity of *M. tuberculosis* containing phagosomes and may contribute to host control of infection.

The ligand-receptor pair CD30 ligand (*Tnfsf8)*, also known as CD153, and CD30 (*Tnfrsf8*) has generated excitement as a potentially important IFN-γ-independent signaling pathway during *M. tuberculosis* infection. In humans, higher frequencies of *M. tuberculosis* specific CD153+ CD4 T cells correlate with a lower lung bacterial load (46), and *Tnfsf8^-/-^* mice have earlier mortality than wild-type following infection (25). However, data point to a regulatory, rather than bactericidal, role for CD153, particularly in the lung. *Tnfsf8* expression is mostly dispensable for control of infection outside of the lungs, and the overabundance of IFN-γ-producing CD4 T cells observed in the lungs of *Tnfsf8^-/-^* mice following infection has been shown to drive immunopathology (13, 46). In a similar manner, reconstitution of *Rag1^-/-^* mice with CD4 T cells that overexpress IFN-γ induces pulmonary pathology and exacerbates bacterial burden in the lung while controlling infection outside of the lungs (13). *Rag1^-/-^* mice reconstituted with *Tnfsf8^-/-^* CD4 T cells have decreased survival compared to wild-type, but lowering the frequency of IFN-γ-producing CD4 T cells in the lung by reconstitution with a 1:1 ratio of *Tnfsf8^-/-^* and *Ifng^-/-^* CD4 T cells rescues this phenotype (25). We show that BMDMs have low expression of *Tnfrsf8* at baseline and after CD4 T cell co-culture, and that BMDMs deficient in *Ifngr* and *Tnfrsf8* have no loss of CD4 T cell-mediated control compared to macrophages lacking *Ifngr* alone. Collectively, these data point to a model where CD153 negatively regulates CD4 T cell IFN-γ production to control immune pathology, particularly in the lung, rather than signaling through CD30 on infected macrophages to induce cell-intrinsic control of bacterial replication.

Mouse and human T cells make GM-CSF during *M. tuberculosis* infection (15), and *Csf2^-/-^* mice have a higher lung bacterial burden and succumb more quickly to disease than wild-type (16). Chimeric mouse experiments with *Csf2^-/-^* and wild-type mice show the GM-CSF production is particularly important for control of infection in the lung, with no effect in the spleen (15). Furthermore, adoptive transfer of wild-type or *Csf2^-/-^* CD4 T cells into wild-type or *Csf2^-/-^* mice shows that CD4 T cell-derived GM-CSF is important *in vivo* only in the absence of non-hematopoietic GM-CSF (15). These *in vivo* experiments are insufficient to conclude that CD4 T cell-derived GM-CSF acts directly on infected macrophages and are complicated by the immunoregulatory roles of GM-CSF, including the fact that *Csf2^-/-^* mice and humans with deficiencies in GM-CSF Receptor signaling have a defect in alveolar macrophage development (47, 48). Our finding that GM-CSF is required for CD4 T cell-mediated IFN-γ-independent control is the first evidence that CD4 T cell-derived GM-CSF participates in cell-intrinsic control of *M. tuberculosis*.

While rGM-CSF is sufficient to restrict *M. tuberculosis* in peritoneal macrophages, it is necessary but not sufficient for control in BMDMs and, surprisingly, does not rescue the lack of IFN-γ-independent control seen in co-cultures with *Csf2^-/-^* CD4 T cells. One explanation for this discrepancy may be that *Csf2^-/-^* CD4 T cells lack an additional effector produced by wild-type T cells that is required for GM-CSF-mediated control in BMDMs. This hypothesis is supported by literature showing that GM-CSF is important for the development, activation, and function of CD4 T cells (49). *Csf2^-/-^* mice have a diminished Th1 response compared to wild-type (50), which partially accounts for the increased susceptibility of *Csf2^-/-^* mice to *M. tuberculosis* infection (16). In anti-tumor immunity, GM-CSF treatment increases the frequency of tumor-specific Th1 T cells (51), and, in experimental autoimmune thyroiditis, GM-CSF treatment has been shown to both enhance IL-6–dependent Th17 cell responses and increase the number of immunoregulatory CD4+CD25+ T cells (52, 53). Alternatively, there may be differences in the ability of CD4 T cell-derived and recombinant GM-CSF to effectively signal to BMDMs. *In vivo*, GM-CSF is subject to substantial post-translation modifications—primarily N- and O-glycosylation—with a molecular weight ranging from 14 to 32 kDa (54), and glycosylation of GM-CSF is required for effective signaling and superoxide induction in neutrophils (55). Western blot analysis of the supernatant of *in vitro* differentiated Th1s revealed at least seven distinct species of GM-CSF, including a heavily glycosylated 32 kDa variant, compared to the single 14 kDa band for non-glycosylated recombinant GM-CSF (Fig S5E). Furthermore, the finding that HIF-1α is required for control in BMDMs by *Ifng^-/-^* CD4 T cells and for control in peritoneal macrophages by rGM-CSF suggests similar mechanisms of action and argues that GM-CSF, rather than a second signal, from *Ifng^-/-^* CD4 T cells is the critical factor for IFN-γ-independent control. Further studies that delineate the role of GM-CSF in CD4 T cell development and macrophage activation will aid the development of vaccines capable of eliciting *M. tuberculosis*-specific GM-CSF-producing CD4 T cells and may expediate an end to the global TB pandemic.

## Methods

### Ethics statement

All procedures involving the use of mice were approved by the University of California, Berkeley Institutional Animal Care and Use Committee (protocol R353-1113B). All protocols conform to federal regulations and the National Research Council’s *Guide for the Care and Use of Laboratory Animals*.

### Reagents

Recombinant mouse IFN-γ was obtained from R&D Systems (485-MI) and used at 6.25 ng/mL. Recombinant mouse GM-CSF expressed in HEK293 cells was obtained from Sino Biological (51048-MNAH) and used at 10 ng/mL. Proteinase K was obtained from Millipore Sigma (RPROTK-RO); 20 mg/mL stocks in ddH_2_O were diluted to a working concentration of 200 μg/mL in cell culture supernatant. 2-Deoxy-D-glucose (2-DG) was obtained from Sigma Aldrich (D8375) and used at 0.32 mM. Supernatant nitrite was measured by the Griess test as a proxy for NO production by mixing a solution of .1% napthylethylenediamine, 1% sulfanilamide, and 2% phosphoric acid 1:1 with supernatant and measuring absorbance at 546nm.

### Mice

C57BL/6 (wild-type, strain 000664), *B6.129S7-Ifng^tm1Ts^/J (Ifng^-/-^*, strain 002287), B6.129P2-*Nos2^tm1Lau^/J (Nos2^-/-^*, strain 002609), and *B6.129S-Csf2^tm1Mlg^/J (Csf2^-/-^*, strain 026812) were purchased from The Jackson Laboratory and bred in-house. *B6.129-Hif1a^tm3Rsjo^/J (Hif1a^fl/fl^*, strain 007561) and *B6.129P2-Lyz2^tm1(cre)Ifo^/J* (LysMcre, strain 004781) mice were obtained from The Jackson Laboratory, crossed to generate *Hif1a^fl/fl^*, LysMcre^+/+^ (referred to here as *Hif1a^-/-^*) mice and bred in-house. *Ifngr1^-/-^ (Ifngr^-/-^*) mice were provided by D. Raulet. *Tnfrsf1a^-/-^/Tnfrsf1b^-/-^ (Tnfr1/2^-/-^*) mice were provided by G. Barton (56). *Ifnar^-/-^/Ifngr1^-/-^ (Ifnar^-/-^/Ifngr^-/-^*) mice were provided by M. Welch (57). *Csf2rb^-/-^* mice were provided by S. Shin (58). C7 TCR tg.CD90.1 (C7 Tg) mice, which express a T cell receptor specific for the *M. tuberculosis* antigen ESAT-6, were provided by K. Urdahl and have been described previously (59, 60). H11^Cas9^ CRISPR/Cas9 knock-in (Cas9 Tg) mice, which constitutively express CRISPR associated protein 9 (Cas9), were provided by R. Vance and were crossed to *Ifngr^-/-^* mice to generate *Ifngr^-/-^* Cas9 Tg mice.

### Cell culture

Bone-marrow was obtained from wild-type, *Nos2^-/-^, Hif1a^-/-^, Ifngr^-/-^, Tnfr1/2^-/-^, Ifnar^-/-^/Ifngr^-/-^*, and *Csf2rb^-/-^* mice and cultured in DMEM with 10% FBS, 2 mM L-glutamine, and 10% supernatant from 3T3–M-CSF cells (BMDM media) for 6 d with feeding on day 3 to generate bone marrow-derived macrophages (BMDMs). Peritoneal macrophages were obtained from wild-type and *Hif1a^-/-^* mice by 5 mL ice-cold PBS lavage and cultured in RPMI with 10% FBS, 2 mM L-glutamine and 35 μg/mL kanamycin (peritoneal macrophage media) for 24 h to allow for macrophage adherence.

### CRISPR/Cas9 gene targeting

Guide sequences targeting *Tnfrsf8* were selected from the murine Brie guide library and cloned into the pLentiGuidePuro backbone from Addgene (52963). Bone marrow was obtained from *Ifngr^-/-^*/Cas9 Tg mice, treated with ACK lysis buffer, and cultured in BMDM media. Lenti-X 293T cells from Takara Bio (632180) were co-transfected with psPAX2, pMD2.G, and pLentiGuidePuro containing target guide sequences using Lipofectamine 2000 from Invitrogen (11668019) according to the manufacturer’s protocol. The next day, 293T media was replaced with BMDM media for 24 hours to collect lentivirus, and *Ifng^-/-^*/Cas9 Tg BMDMs were then transduced using 293T supernatant. On day 5, 4 μg/mL puromycin was added to select for a polyclonal population of cells. On day 12, *Ifngr^-/-^/Tnfrsf8^-/-^* double-knockout BMDMs were harvested and infected with *M. tuberculosis* as described above. Targeting efficiency was determined by sequencing targeted genomic sites and analyzing population level genome editing using TIDE analysis (https://tide.deskgen.com/) (61). *Tnfrsf8* was targeted in two independent experiments, each with three targeting guides. Data shown in Fig 4 are representative of results with gRNA 5’-AGACCTCAGCCACTACCCAG-3’ which had genome editing efficacy of 91.3–91.9%.

### Bacterial cultures

Frozen stocks of low passage *M. tuberculosis* Erdman were grown to log phase over 5 d at 37°C in Middlebrook 7H9 media with 10% albumin-dextrose-saline, 0.5% glycerol, and 0.05% Tween-80. Δ*EccC M. tuberculosis* Erdman and *M. tuberculosis* Erdman-lux strain expressing luciferase from the *luxCDABE* operon have been described previously and were cultured as described above (62, 63). *M. tuberculosis* Erdman-mCherry was generated by D. Kotov and R. Vance by transforming wild-type Erdman with pMSP12::mCherry, a gift from Lalita Ramakrishnan (Addgene plasmid # 30169; http://n2t.net/addgene:30169; RRID:Addgene_30169) and was cultured as described above.

### *In vitro* infections

BMDMs were plated in 24-well or 96-well plates at 3 x 10^5^ or 5 x 10^4^ cells/well, respectively, allowed to adhere for 48 hours, and infected in DMEM with 5% horse serum and 5% FBS by 4-hour phagocytosis or 10 min spinfection at 300 rcf. Unless otherwise indicated, BMDMs were infected at a multiplicity of infection of 1. Peritoneal macrophages were plated in 12-well, 24-well or 96-well plates at 1.65 x 10^6^, 1.1 x 10^6^ or 1.5 x 10^5^ cells/well, respectively, cultured for 24 hours to allow for macrophage adherence, washed with PBS, and infected in peritoneal macrophage media with 5% horse serum by 10 min spinfection at 300 rcf. After infection, cells were washed once with PBS before replacing with the appropriate media. CFU/well was determined at the indicated time points by washing infected cells with PBS, lysing in sterile-filtered distilled water with 0.5% Triton-X100 for 10 min at 37°C, diluting in PBS prepared with 0.05% Tween 80, plating onto 7H10 plates supplemented with 0.5% glycerol and 10% OADC (Middlebrook), and incubating for 21 d at 37°C. CFU is reported as fold-change over inoculum (calculated by plating t=0 CFU immediately after phagocytosis or spinfection), or as CFU/well, referring to the total number of bacteria per well of the assay plate. Luminescence emission of *M. tuberculosis* Erdman-lux was measured at 470 nm with a Spectra-L plate reader (Molecular Devices, San Jose, CA).

### Lung-derived CD4 T cell co-cultures

CD4+ cells were isolated from the lungs of 7–12-week-old wild-type, *Ifng^-/-^, Nos2^-/-^* or *Csf2^-/-^* mice 21 – 28 d after ~500 CFU aerosol infection with *M. tuberculosis* Erdman using mouse CD4 (L3T4) MicroBeads from Miltenyi Biotec (130-117-043) according to the manufacturer’s protocol, resuspended in RPMI with 10% FBS, 2 mM L-glutamine, 1 mM Sodium pyruvate, 1 mM NEAA, 20 mM HEPES, 55 μM 2-mercaptoethanol and 35 μg/mL kanamycin (T cell media) supplemented with 10% supernatant from 3T3–M-CSF cells and added to infected BMDMs in co-culture or in transwells. Unless indicated, CD4 T cells were added at a 5:1 T cell:BMDM ratio. In these experiments, T cell media supplemented with 10% supernatant from 3T3–M-CSF cells was used for untreated and IFN-γ-treated controls.

### *In vitro* T cell differentiation

CD4+ cells were isolated from C7 Tg mouse spleens using mouse CD4 (L3T4) MicroBeads from Miltenyi Biotec (130-117-043) according to the manufacturer’s protocol and cultured in T cell media at 2 x 10^6^ cells/mL with 1 μM ESAT-6 peptide pool (NIH BEI Resources Repository catalog no. NR-34824) and 2 x 10^6^ cells/mL irradiated wild-type splenocytes (3200 rads). To differentiate Th1s, 10 U/mL IL-2 from R&D Systems (1150-ML), 5 U/mL IL-12 p70 from PeproTech (210-12), and 1 μg/mL rat anti-mouse IL-4 from R&D Systems (MAB404100) was added; for Th17.1s, 2 ng/mL TGF-β from Invitrogen (14834262), 20 ng/mL IL-6 from PeproTech (216-16), 2.5 μg/mL rat anti-mouse IFN-γ from Biolegend (505801) and 1 μg/mL anti-IL-4 was added; and for Th2s, 10 U/mL IL-2, 10 ng/mL IL-4 from R&D Systems (404-ML), and 3 μg/mL anti-IFN-γ was added. On day 3, cells were collected and replated at 2 x 10^6^ cells/mL with 10 U/mL IL-2 (Th1s), 2 ng/mL TGF-β, 10 ng/mL IL-6, 30 ng/mL IL-1β from R&D Systems (401-ML), 0.5 μg/mL anti-IFN-γ, and 0.33 μg/mL anti-IL-4 (Th17.1s), or 10 U/mL IL-2, 10 ng/mL IL-4, and 1 μg/mL anti-IFN-γ (Th2s). Differentiated T cells were harvested on day 5 and used in co-culture at a 5:1 T cell:BMDM ratio or were replated at 2 x 10^6^ cells/mL in T cell media and stimulated overnight with 100 U/mL IL-2 (Th1s), 5 ng/mL IL-1β and 5 ng/mL IL-23 from Invitrogen (14823163) (Th17.1s) or 100 U/mL IL-2 and 100 ng/mL IL-4 (Th2s) in plates coated with 10 μg/mL Armenian hamster anti-mouse CD3ε (BE0001-1) and 2 μg/mL Syrian hamster anti-mouse CD28 (BE0015-1) from Bio X Cell to generate Th1, Th17.1, or Th2 supernatants. On day 6, T cell supernatants were collected and supplemented with 10% supernatant from 3T3–M-CSF cells before addition to BMDMs. In these experiments, T cell media supplemented with 10% supernatant from 3T3–M-CSF cells was used for untreated and IFN-γ-treated controls.

### Cytokine profiling

Differentiated Th1 and Th17.1 cells were harvested on day 5, replated at 2 x 10^6^ cells/mL in Opti-MEM media (Gibco) plus 2 mM L-glutamine, 1 mM Sodium pyruvate, 1 mM NEAA, 20 mM HEPES and 55 μM 2-mercaptoethanol (Opti-MEM T cell media) and stimulated overnight as described above. On day 6, T cell supernatants were collected, and cytokine profiling was performed by the Stanford Human Immune Monitoring Center (Stanford, CA) using a mouse 48-plex immunoassay (Procarta).

### RNA sequencing

For RNA-seq on BMDMs after lung CD4 T cell co-culture (Fig 2), four independent experiments were performed. For each experiment, BMDMs were seeded in 24-well plates, infected as described, and co-cultured with lung CD4 T cells isolated as described. At 24 hours postinfection, CD4 T cells were separated from BMDMs using CD4 (L3T4) Dynabeads from Invitrogen (11445D) according to the manufacturer’s protocol, and BMDMs were lysed in 1 mL TRIzol (Invitrogen). Total RNA was extracted using chloroform and 2 mL Heavy Phase Lock Gel tubes from QuantaBio (2302830), and the aqueous layer was further purified using RNeasy Mini spin columns from Qiagen (74104). RNA quality was determined using an Agilent 2100 Bioanalyzer and RNA concentration was determined using the Qubit Quantitation Platform at the Functional Genomics Laboratory of The California Institute for Quantitative Biosciences (University of California, Berkeley). RNA-seq libraries were prepared by the DNA Technologies and Expression Analysis Core (University of California, Davis) and differential gene expression was analyzed by the Genome Center and Bioinformatics Core Facility (University of California, Davis).

### HIF-1α western blot

BMDMs infected for 12 hours at a multiplicity of infection of 5 as described were washed with ice cold PBS, lysed in 1X SDS-PAGE buffer on ice and heat sterilized for 20 min at 100°C for removal from the BSL3 facility. Total protein lysate was analyzed by SDS-PAGE using precast 4-20% Criterion TGX protein gels from Bio-Rad Laboratories (5671093), rabbit mAb to HIF-1α (D2U3T) from Cell Signaling Technology (14179S), and an HRP-conjugated secondary antibody. Blots were developed with Western Lightning Plus-ECL chemiluminescence substrate from PerkinElmer (NEL105001EA) and a ChemiDoc MP system from Bio-Rad Laboratories.

### Microscopy

For immunoflourescence microscopy, BMDMs were plated at 3 x 10^4^ cells/well in black, clear bottom 96-well plates and infected with *M. tuberculosis* Erdman-mCherry as described above at a multiplicity of infection of 2. Eight hours postinfection, cells were fixed in 4% paraformaldehyde, washed with PBS and immunostained for polyubiquitin (FK2, Millipore Sigma 04-263, 1:400 dilution) or Galectin-3 (B2C10, sc-32790, 1:50 dilution) at 4C overnight, followed by secondary antibody staining at room temperature for 30 minutes. For microscopy to detect lipid droplets and intracellular lipid inclusions, BMDMs or peritoneal macrophages were plated at 5 x 10^4^ or 1.5 x 10^5^ cells/well, respectively, in black, clear bottom 96-well plates and infected with *M. tuberculosis* Erdman-mCherry as described above at a multiplicity of infection of 4 (BMDMs) or 2 (peritoneal macrophages). At the indicated time points, cells were fixed in 4% paraformaldehyde, washed with PBS and stained for neutral lipids with BODIPY 493/503 from Invitrogen (D3922) at 1 μg/mL and with DAPI. All imaging was done on a Perkin Elmer Opera Phenix Automated Microscopy System.

### Statistical Analysis

Analysis of statistical significance was performed using GraphPad Prism 8 (GraphPad, La Jolla, CA).

## Acknowledgements

*Csf2rb^-/-^, Ifngr^-/-^, Ifngr^-/-^/Ifnar^-/-^ Nos2^-/-^, Tnfrsf1a^-/-^/Tnfrsf1b^-/-^*, C7 Tg, and Cas9 Tg mice were a gift from S. Shin, D. Raulet, M. Welch, R. Vance, G. Barton, K. Urdahl and R. Vance, respectively. We thank T. Burke, D. Kotov, K. Lien, K. Magee, S. Margolis, X. Nguyenla, C. Nicolai, A. Olive, A. Roberts, and L. Shallow for technical assistance. We thank Y. Rosenberg-Hasson and the Human Immune Monitoring Center (Stanford University) for cytokine profiling, the Functional Genomics Laboratory (UC Berkeley) for RNA sample quality check, and L. Froenicke at the DNA Technologies and Expression Analysis Core (UC Davis) and at the Genome Center and Bioinformatics Core Facility (UC Davis) for RNAseq and data analysis. Finally, we thank G. Barton, M. DuPage, R. Vance and members of the Cox and Stanley labs for helpful discussions. This work was supported by NSF Graduate Research Fellowship DGE-1752814 to E.V.D. and funding from the National Institute of Allergy and Infectious Diseases (1R01AI143722 to S.A.S and P01 AI063302 and U19 AI106754 to J.S.C. and S.A.S).

## Author Contributions

Conceptualization, Methodology, and Analysis: EVD and SAS

Investigation: EVD, HMM, DMF, JPB, LHM, and SR

Writing – original draft: EVD

Writing – review & editing: EVD, HMM, DMF, JSC and SAS

Funding Acquisition: EVD, JSC and SAS

Supervision: JSC and SAS

## Disclosures

The authors declare no competing interests.

## Extended Methods

### Reagents

GM-CSF expressed in *E. coli* was obtained from R&D Systems (415-ML) and GM-CSF expressed in Chinese hamster ovary cells was obtained from MedChemExpress (HY-P7069) and used at 10 ng/mL. Armenian hamster anti-mouse CD40 (102907) was obtained from Biolegend and used at 20 μg/mL. Glucose uptake was determined using a glucose (HK) assay kit from Sigma-Aldrich (GAHK20) according to the manufacturer’s protocol.

### Transcription factor prediction

oPOSSUM has been described previously (1) and was used to predict macrophage transcription factors regulated by CD4 T cells during IFN-γ independent control of infection. oPOSSUM analysis was run on all genes in the RNAseq data found to be expressed at a higher level in *Ifngr^-/-^* BMDMs after co-culture with lung-derived CD4 T cells compared to UT.

### GM-CSF ELISA

GM-CSF concentration in cell culture supernatant was determined using the Mouse GM-CSF Quantikine ELISA kit from R&D Systems (MGM00) according to the manufacturer’s protocol.

### GM-CSF western blot

C7 Th1 supernatants were generated in OptiMEM T cell media as described. Protein from 1 mL supernatant was concentrated by TCA precipitation and compared to a dose response of recombinant GM-CSF by SDS-PAGE using precast 4-20% Criterion TGX protein gels from Bio-Rad Laboratories (5671093), rabbit polyclonal Ab to mouse GM-CSF from Abcam (ab9741), and an HRP-conjugated secondary antibody. Blots were developed as described in the main text.

---

**Figure S1.**
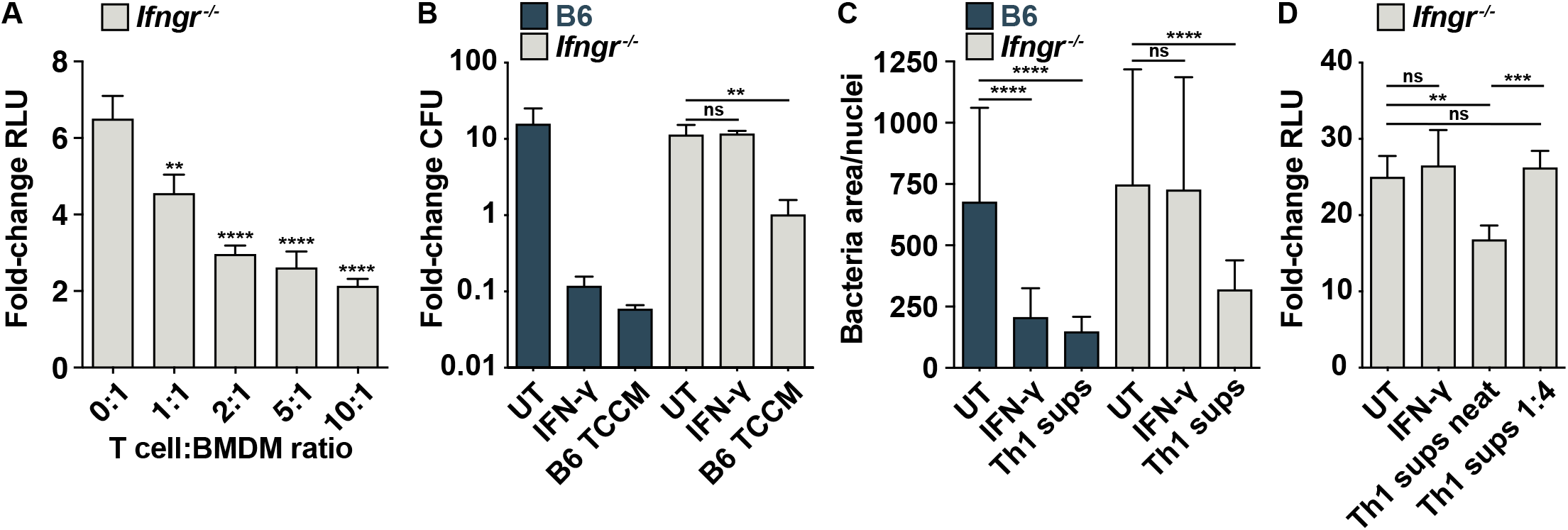
Related to Fig 1, lung-derived and *in vitro*-differentiated CD4 T cells control *M. tuberculosis* growth independent of IFN-γ. (**A**) RLU fold-change at d 4 postinfection for *Ifngr^-/-^* BMDMs co-cultured with the indicated ratios of lung-derived wild-type CD4 T cells to BMDMs, p-values relative to UT. (**B**) CFU fold-change at d 5 postinfection for wild-type and *Ifngr^-/-^* BMDMs treated with conditioned media from lung-derived wild-type CD4 T cells (TCCM). (**C**) Microscopy for bacterial replication at d 3 postinfection for wild-type and *Ifngr^-/-^* BMDMs treated with Th1 supernatants. (**D**) RLU fold-change at d 4 postinfection for *Ifngr^-/-^* BMDMs treated with neat or diluted Th1 supernatants. Figures are representative of two (A), (D) or at least three (B)-(C) independent experiments. Error bars are SD from four replicate wells (A)-(B), (D) or 48 images from 4 replicate wells (C), **p<0.01, ***p<0.001, ****p<0.0001 by unpaired t-test.

**Figure S2.**
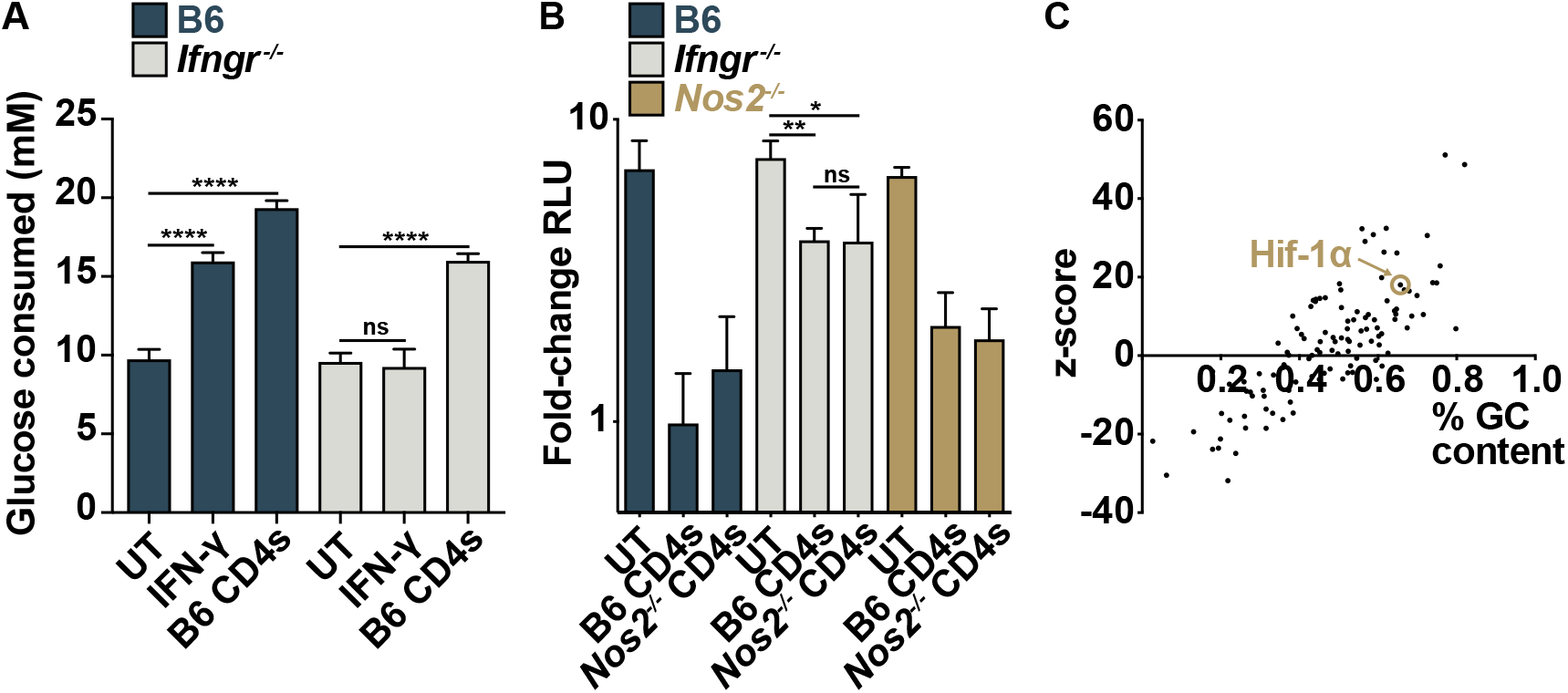
Related to Fig 2, CD4 T cells induce an IFN-γ- and NO-independent increase in glycolytic gene expression. (**A**) Glucose consumption at 48 h postinfection for wild-type and *Ifngr^-/-^* BMDMs co-cultured with lung-derived wild-type CD4 T cells. (**B**) RLU fold-change at d 6 postinfection for wild-type, *Ifngr^-/-^* and *Nos2^-/-^* BMDMs co-cultured with a 10:1 ratio of lung-derived wild-type or *Nos2^-/-^* CD4 T cells. (**C**) oPOSSUM bioinformatic prediction of transcription factors responsible for regulation of all genes found by RNA sequencing to be expressed at a higher level in *Ifngr^-/-^* BMDMs after co-culture with lung-derived CD4 T cells compared to UT. Figures represent data from three independent experiments (C) or are representative of two independent experiments (A)-(B). Error bars are SD from four replicate wells, *p<0.05, **p<0.01, ****p<0.0001 by unpaired t-test.

**Figure S3.**
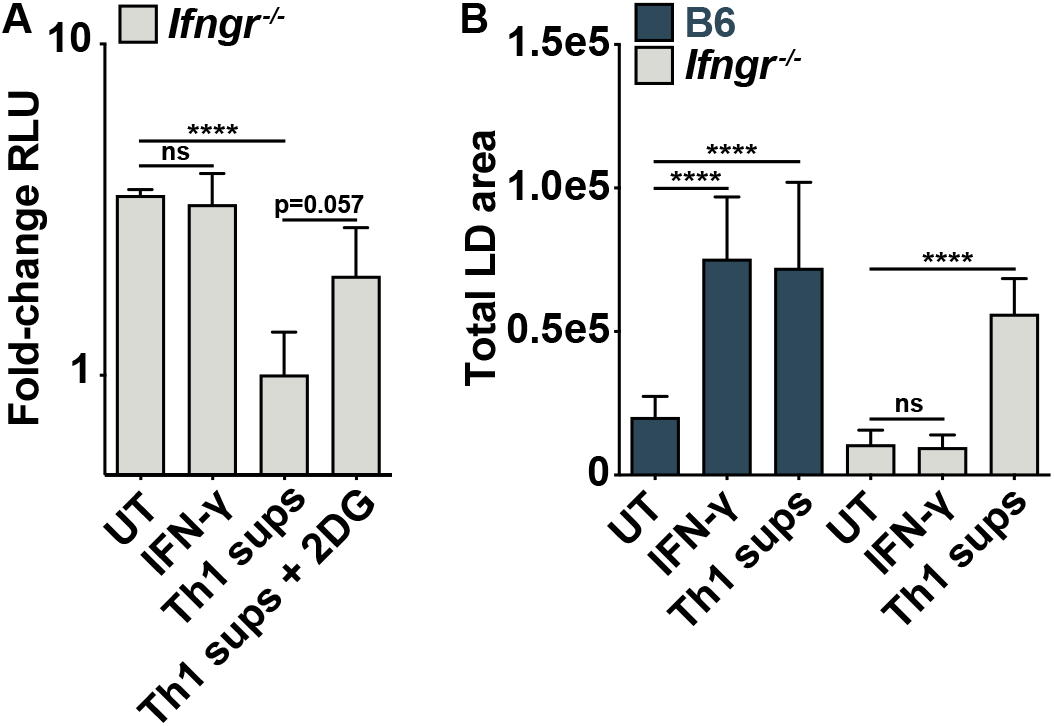
Related to Fig 3, IFN-γ-independent control of *M. tuberculosis* requires the macrophage transcription factor HIF-1α. (**A**) RLU fold-change at d 4 postinfection for wild-type and *Ifngr^-/-^* BMDMs treated with Th1 supernatants and 2-DG. (**B**) Quantification of (Fig 3C-F) for total area of LDs in the imaging field. Figures are representative of two (A) or three (B) independent experiments. Error bars are SD from four replicate wells (A) or 48 images from four replicate wells (B), ****p<0.0001 by unpaired t-test.

**Figure S4.**
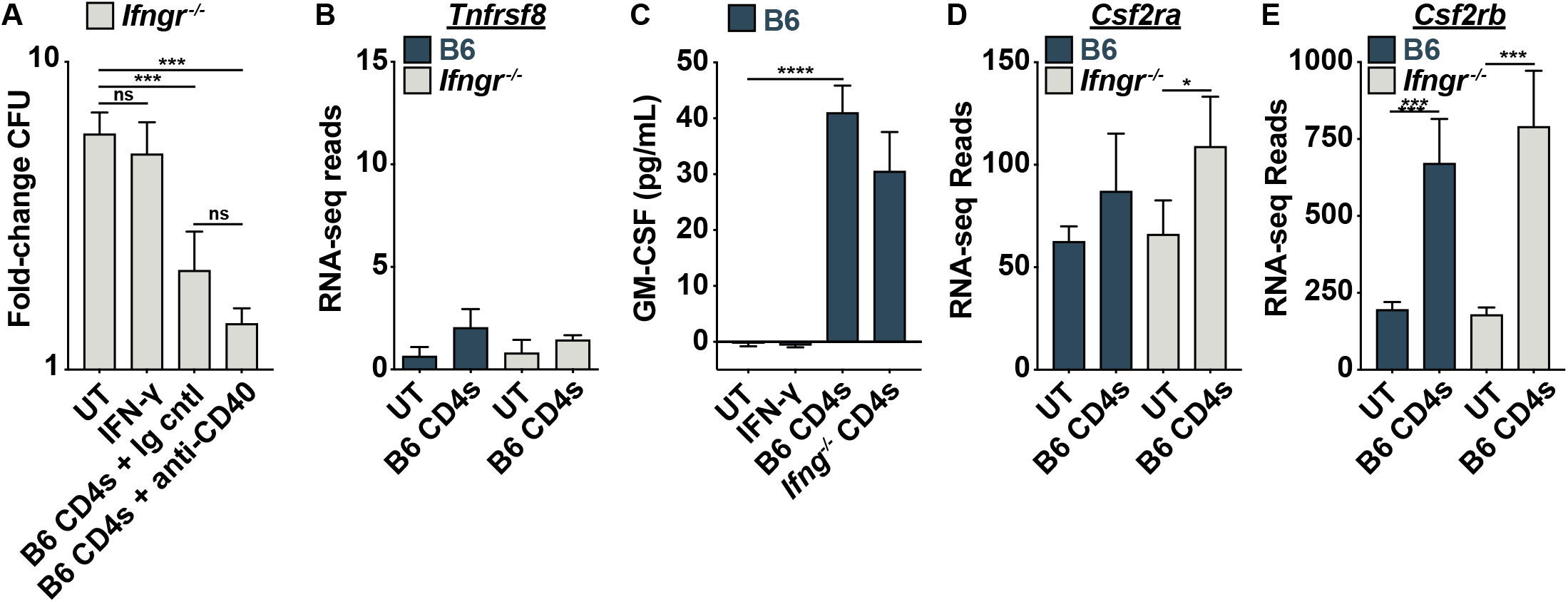
Related to Fig 4, IFN-γ-independent control of *M. tuberculosis* requires CD4 T cell-derived GM-CSF. (**A**) CFU fold-change at d 5 postinfection for *Ifngr^-/-^* BMDMs co-cultured with lung-derived wild-type CD4 T cells and treated with anti-CD40. (**B**) RNA-seq reads of *Tnfrsf8* at 24 h postinfection in wild-type and *Ifngr^-/-^* BMDMs co-cultured with lung-derived CD4 T cells. (**C**) ELISA for GM-CSF concentration in lung CD4 co-culture supernatants at d 2 postinfection. (**D**)-(**E**) RNA-seq reads of (D) *Csf2ra* and (E) *Csf2rb* at 24 h postinfection in wild-type and *Ifngr^-/-^* BMDMs co-cultured with lung-derived CD4 T cells. Figures show three biological replicates (B), (D)-(E) or are representative of three independent experiments (A), (C). Error bars are SD from three independent experiments (B), (D)-(E) or four replicate wells (A), (C), *p<0.05, ***p<0.001, ****p<0.0001 by unpaired t-test.

**Figure S5.**
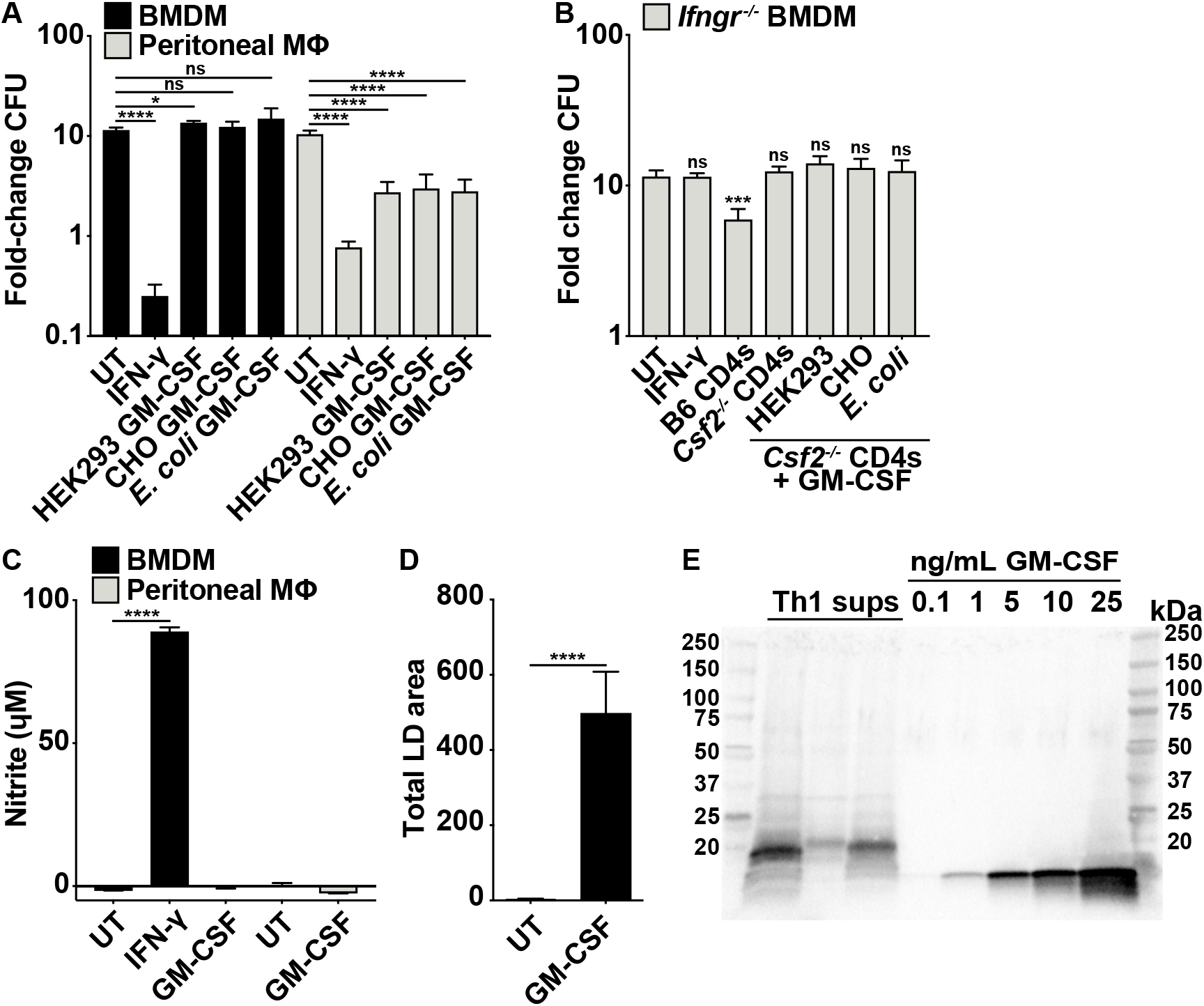
Related to Fig 5, HIF-1α-dependent restriction of *M. tuberculosis* by GM-CSF occurs only in peritoneal macrophages. (**A**) CFU fold-change at d 4 postinfection for BMDMs and peritoneal macrophages treated with GM-CSF derived from the indicated sources. (**B**) CFU fold-change at d 5 postinfection for *Ifngr^-/-^* BMDMs co-cultured with wild-type or *Csf2^-/-^* lung CD4 T cells and treated with GM-CSF derived from the indicated sources, p-values relative to UT. (**C**) Griess assay at 24 h postinfection for BMDMs and peritoneal macrophages treated with GM-CSF. (**D**) Quantification of (Fig 5C-D) for total area of LDs in the imaging field of peritoneal macrophages treated with GM-CSF. (**E**) Western blot for GM-CSF comparing a dose response of recombinant GM-CSF to three biological replicates of 1mL TCA-precipitated C7 Th1 supernatants. Figures are representative of two (B)-(C) or at least three (A), (D)-(E) independent experiments. Error bars are SD from four replicate wells (A)-(C) or 48 images from four replicate wells (D), *p<0.05, ***p<0.001, ****p<0.0001 by unpaired t-test.

## Notes

### Competing Interest Statement

The authors have declared no competing interest.

